# Factors Promoting Lipopolysaccharide Uptake by Synthetic Lipid Droplets

**DOI:** 10.1101/2024.10.19.619182

**Authors:** Assame Arnob, Anirudh Gairola, Hannah Clayton, Arul Jayaraman, Hung-Jen Wu

## Abstract

Lipoproteins are essential in removing lipopolysaccharide (LPS) from blood during bacterial inflammation. The physicochemical properties of lipoproteins and environmental factors can impact LPS uptake. In this work, synthetic lipid droplets containing triglycerides, cholesterols, and phospholipids, were prepared to mimic lipoproteins. The physicochemical properties of these lipid droplets, such as charges, sizes, and lipid compositions, were altered to understand the underlying factors affecting LPS uptake. The amphiphilic LPS could spontaneously adsorb on the surface of lipid droplets without lipopolysaccharide binding protein (LBP); however, the presence of LBP can increase LPS uptake. The positively charged lipid droplets also enhance the uptake of negatively charged LPS. Most interestingly, the LPS uptake highly depends on the concentrations of Ca^2+^ near the physiological conditions, but the impact of Mg^2+^ ions was not significant. The increase of Ca^2+^ ions can improve LPS uptake by lipid droplets; this result suggested that Ca^2+^ may play an essential role in LPS clearance. Since septic shock patients typically suffer from hypocalcemia and low levels of lipoproteins, the supplementation of Ca^2+^ ions along with synthetic lipoproteins may be a potential treatment for severe sepsis.

## 1. INTRODUCTION

Lipopolysaccharide (LPS), also known as endotoxin, is a vital component of the outer cell membrane of gram-negative bacteria.^1^ LPS consists of a hydrophobic lipid moiety, called lipid A, along with a distal polysaccharide known as O-antigen, and an oligosaccharide core (**Figure 1a**). LPS plays an important role in inducing the toxicity elicited by gram-negative bacteria. LPS binds to a series of pattern recognition receptors, including the toll-like receptor 4 (TLR4), and triggers an inflammatory response leading to the production of pro-inflammatory cytokines. Overproduction of pro-inflammatory molecules can lead to septic shock, resulting in fatal clinical consequences. Neutralization of LPS is one of the therapeutic approaches to reduce mortality in septic patients.^2^ For example, hemoperfusion and anti-LPS antibodies have been developed to treat septic shock, but the clinical trials showed inconclusive results.^3, 4^

**Figure 1:**
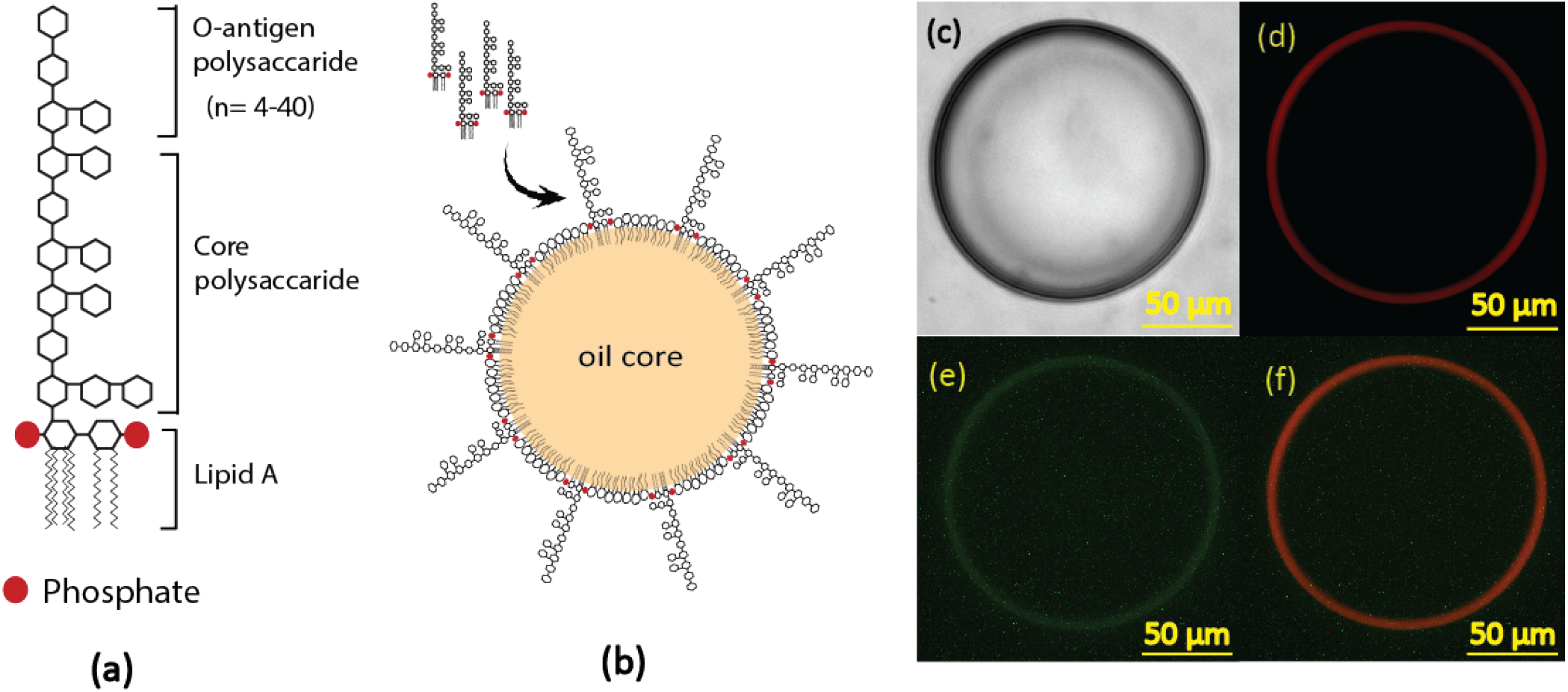
(a) Structure of lipopolysaccharide (LPS), red circles represent negatively charged phosphates linked to lipid A, (b) Schematic for the possible LPS adsorption onto lipid droplets, (c) Giant lipid droplet under brightfield, (d) Epifluorescence microscope image of the lipid droplet labeled with Texas Red-DHPE on the droplet surface, (e) FITC labeled LPS (FITC-LPS) stays on the surface of the lipid droplet, (f) The merger of Texas Red and FITC channels.

Another LPS neutralization strategy is using natural or synthetic lipoproteins to clear LPS from the bloodstream. It is known that lipoproteins play an important role in clearing endotoxins.^5^ Septic patients typically suffer from lower levels of lipoproteins, which could lead to worse clinical outcomes.^5, 6^ Therefore, administrations of natural or synthetic lipoproteins have been used to inhibit endotoxin-induced mortality and restrict LPS activity. Although preclinical studies demonstrated the efficacy of lipoprotein treatment, results from preclinical and clinical trial are inconsistent.^7-11^ To maximize therapeutic efficacy, it is critical to identify the key parameters influencing LPS-lipoprotein interactions.

Lipoproteins contain a hydrophobic core surrounded by amphiphilic molecules. The hydrophobic core is made up mostly of triglycerides and cholesterol esters, while amphiphilic molecules including, phospholipids, cholesterol, and apolipoproteins, encircle the hydrophobic core.^12^ These lipoproteins are essential for the small intestine’s ability to absorb and transport dietary lipids and move lipids from the liver to peripheral tissues and back to the liver and gut.^13^ Transporting harmful foreign hydrophobic and amphipathic substances, including bacterial endotoxin, from invasion and infection sites is a secondary function. Based on their size, lipid content, and apolipoproteins, plasma lipoproteins are classified into chylomicron remnants, chylomicrons, very low-density lipoproteins (VLDL), low-density lipoproteins (LDL), intermediate-density lipoproteins (IDL), and high-density lipoproteins (HDL). According to recent findings, chylomicron, LDL, and HDL have been shown to increase LPS removal.^14, 15^

The uptake of LPS by lipoproteins removes LPS from circulation, and then LPS attached lipoproteins are broken down in the liver.^13^ Previously, a few physiological parameters impacting LPS clearance have been investigated. For example, a mild elevation of LPS binding protein (LBP) is critical in lowering LPS activity.^14^ LBP is an essential carrier delivering the amphiphilic LPS monomer not only to lymphocyte membranes for TLR4 signaling but also to lipoproteins for LPS clearance.^16^ Increases in LBP concentrations can assist in LPS clearance and reduce immune responses.^17^ Apoprotein, one of the lipoprotein components, is the other variable showing a positive impact on the LPS adsorption onto lipoproteins.^14^ Other physicochemical parameters, however, have not yet been investigated.

In this study, the parameters influencing LPS adsorption to synthetic lipid droplets that mimic lipoproteins were investigated. The synthetic lipid droplets contain a triglyceride core surrounded by phospholipids and cholesterol. The influences of phospholipids, cholesterol compositions, lipid droplet size, LBP concentration, and bivalent cations on LPS adsorption were investigated. Interestingly, calcium ions have a significant impact on LPS adsorption near the physiological concentrations. This finding suggests that calcium supplementation may improve the efficacy of lipoprotein treatment for septic patients.

## 2. RESULTS

### 2.1 LPS adsorption on lipid droplets

Current literature does not fully describe the mechanisms underlying LPS uptake by lipoproteins. As LPS is an amphiphilic molecule, we hypothesized that LPS would adsorb to the surface of the lipid droplet by embedding its lipid tail into the oil core of the lipid droplet. The LPS uptake process was first observed by fluorescence microscopy. For visualization, giant lipid droplets (80∼130 µm in size) labeled with Texas Red-DHPE were fabricated and then incubated with FITC labeled LPS (FITC-LPS). Under fluorescent microscopy, we observed LPS majorly adsorbed on the surface of the lipid droplets (**Figure 1**). To further corroborate the surface adsorption of LPS, the lipid droplet size before and after LPS incubation was measured using dynamic light scattering (DLS) (**Figure S1 & Table S1**). The increase in the hydrodynamic diameter of lipid droplets was observed after LPS adsorption, due to the long polysaccharide chains on LPS molecules.

### 2.2 Influences of lipid droplet size and cholesterol

According to the literature, the insertion of amphiphilic molecules into cell membranes is curvature sensitive.^18^ A high curvature lipid bilayer introduces mismatches to the densely packed phospholipids, thus, exposing hydrophobic areas to the aqueous medium as defects.^19^ Such defects offer sites to recruit amphiphilic molecules. Thus, the lipid droplet sizes may be an important factor for LPS adsorption.

The sizes of the lipid droplets were altered by changing lipid to triglyceride molar ratios. The final lipid droplet sizes were determined by DLS after lipid droplet fabrication (**Table S1)**. For LPS adsorption measurement, the droplets labeled with Texas Red-DHPE were incubated with the FITC-labeled LPS, and the amounts of LPS and droplets were quantified by the fluorescent signals. Because LPS majorly adsorbs on the surface of the lipid droplets, the total surface area of the lipid droplets would influence the amount of LPS adsorption. Therefore, the LPS uptake efficiency is expressed as the ratios of adsorbed LPS molecules to phospholipids in droplets (**Figure 2**). For the lipid droplet (molar composition: 99.25% POPC, 0.25% Texas Red-DHPE, 0.5% biotinyl PE) in 1x PBS buffer, the LPS to phospholipid ratio is 0.29 ± 0.03 µg/µg. The molecular weight of POPC is 760.076 g/mol. According to the literature, the molecular weight of an LPS monomer ranges from 10 to 20 kDa.^20, 21^ Assuming the molecular weight of LPS is 10 kDa, the LPS to phospholipid molar ratio was around 1:45 (i.e., a single LPS molecule was surrounded by 45 phospholipids).

**Figure 2:**
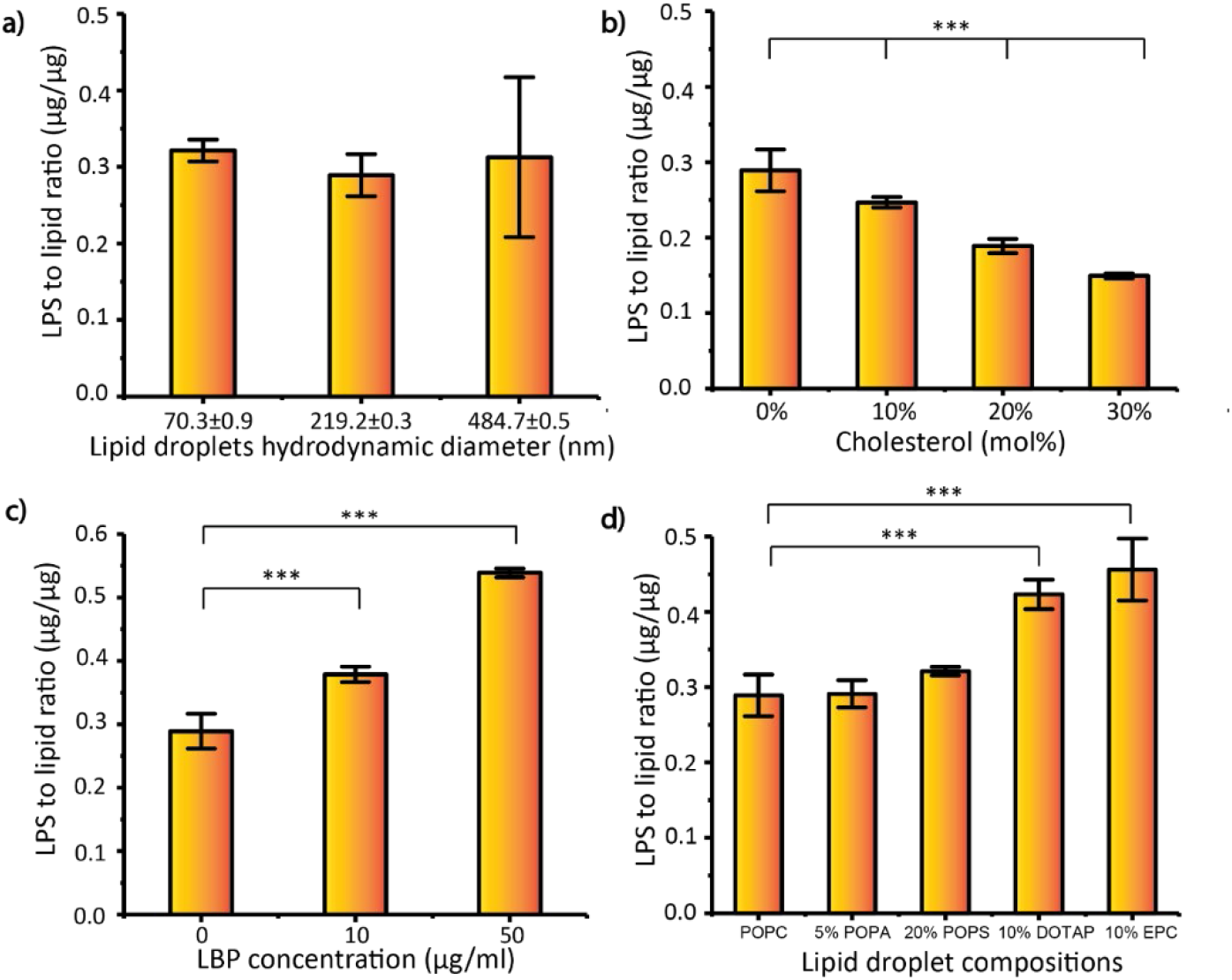
Effect of different parameters on LPS adsorption by lipid droplets. a) effect of lipid droplet size, b) effect of cholesterol, c) effect of LBP concentration, d) effect of charged lipids in lipid droplet composition. For analysis, the data was collected from 3 technical replicates for each run. ***p value<0.001, **p value <0.01.

We prepared lipid droplets in the range of 70 nm to 500 nm in diameter. In this size range, we did not observe a significant impact of droplet size on LPS adsorption (**Figure 2a**). Curvature dependence is also associated with the size of hydrophobic portions in amphiphilic molecules. Prior studies have reported that the interfacial tension becomes significant for lipid droplets with a radius of ∼5 nm when the cross-sectional area per lipid molecule is lower than ∼1.4 nm.^22^ Because the area per LPS molecule is 1.09 nm^2^,^23^ the curvature may play an important role for smaller lipid droplets.

Human lipoproteins contain free cholesterol or cholesterol ester. Chylomicrons can have up to 1-3 wt.% cholesterol ester and free cholesterol^24^, and HDL contains around 28 wt.% cholesterol ester and 6% free cholesterol.^25^ Therefore, we evaluated the influence of cholesterol content by varying cholesterol composition from 0 to 30 mol%. The increase in cholesterol composition in lipid droplets reduced LPS uptake (**Figure 2b**). The lipid droplets containing 30 mol% cholesterol showed approximately 50% of the LPS uptake as compared to the droplet without cholesterol.

### 2.3 Influence of LPS binding protein (LBP) and charged lipids

LBP could disassemble LPS aggregates and deliver the LPS monomer to lymphocyte membranes and lipoproteins.^26^ Studies have shown that the amount of LBP present in the blood of septic patients can be used as an important biomarker for the sepsis process.^27^ Clinical studies have reported that the concentration of LBP in the blood is around 4.1 ± 1.65 µg/ml for healthy individuals and ∼ 31.2 µg/ml (interquartile range, 22.5-47.7 µg/mL) for septic patients.^28^ To observe the influence of LBP, LPS uptake experiments were performed at LBP concentrations ranging from 0 to 50 µg/ml (**Figure 2c**). A positive correlation between LBP and LPS uptake was observed. The increase in LBP concentration improves LPS uptake. Compared to 0 µg/ml LBP concentration (control), the LPS adsorption was almost doubled at 50 µg/ml LBP concentration.

The presence of phosphate groups in lipid A gives the net negative charges of LPS.^29^ The electrostatic interactions among LPS, phospholipids, and environmental ions influence the packing density of LPS on lipid droplets. To investigate the influence between charged lipids and LPS, we mixed cationic and anionic phospholipids with the neutral POPC in the lipid droplet. We introduced 10 mol% DOTAP and 10 mol% EPC to get positively charged lipid droplets. The zeta potential of the droplets was measured by DLS (**Table S1**). The results (**Figure 2d**) show that LPS uptake increased by almost 40% compared to the lipid droplets prepared with neutral POPC. The cationic phospholipids may couple with the negatively charged LPS, facilitating the insertion of LPS into droplet surfaces. In contrast, when we mixed 20 mol% POPS or 5 mol% POPA with POPC in lipid droplets to introduce negatively charged phospholipids on droplet surfaces, the negatively charged phospholipids did not show significant influence on the LPS adsorption.

### 2.4 The influence of bivalent cations (Ca^2+^ and Mg^2+^)

Prior studies have shown that the concentration of Ca^2+^ decreases significantly in septic patients.^30-32^ For a healthy population, the median amount of Ca^2+^ ions is around 2.31 mM^31^, and the Ca^2+^ concentrations in septic patients could be lower than 0.93 mM. In contrast, Mg^2+^ ion concentration does not change much in the blood.^33^ Bivalent cations are known to neutralize and stabilize LPS in the supported lipid bilayers.^29, 34^ To observe the effect of bivalent cations present in the buffer, Ca^2+^ or Mg^2+^ ions were varied from 0 to 150 mM (**Figure 3b**). From 2 to 10 mM of Ca^2+^ ions, the LPS adsorption nearly doubled. This increase in LPS adsorption in the presence of the Ca^2+^ ions may be explained by the thermodynamically favorable insertion of LPS in lipid membranes.^29^ The prior study hypothesized that a Ca^2+^ ions may bridge negatively charged LPS and promotes self-insertion of LPS into a lipid bilayer.^29^ (**Figure 3a**) However, the presence of Mg^2+^ ion in the buffer from 0 to 150 mM did not influence LPS adsorption (**Figure 3b**).

**Figure 3:**
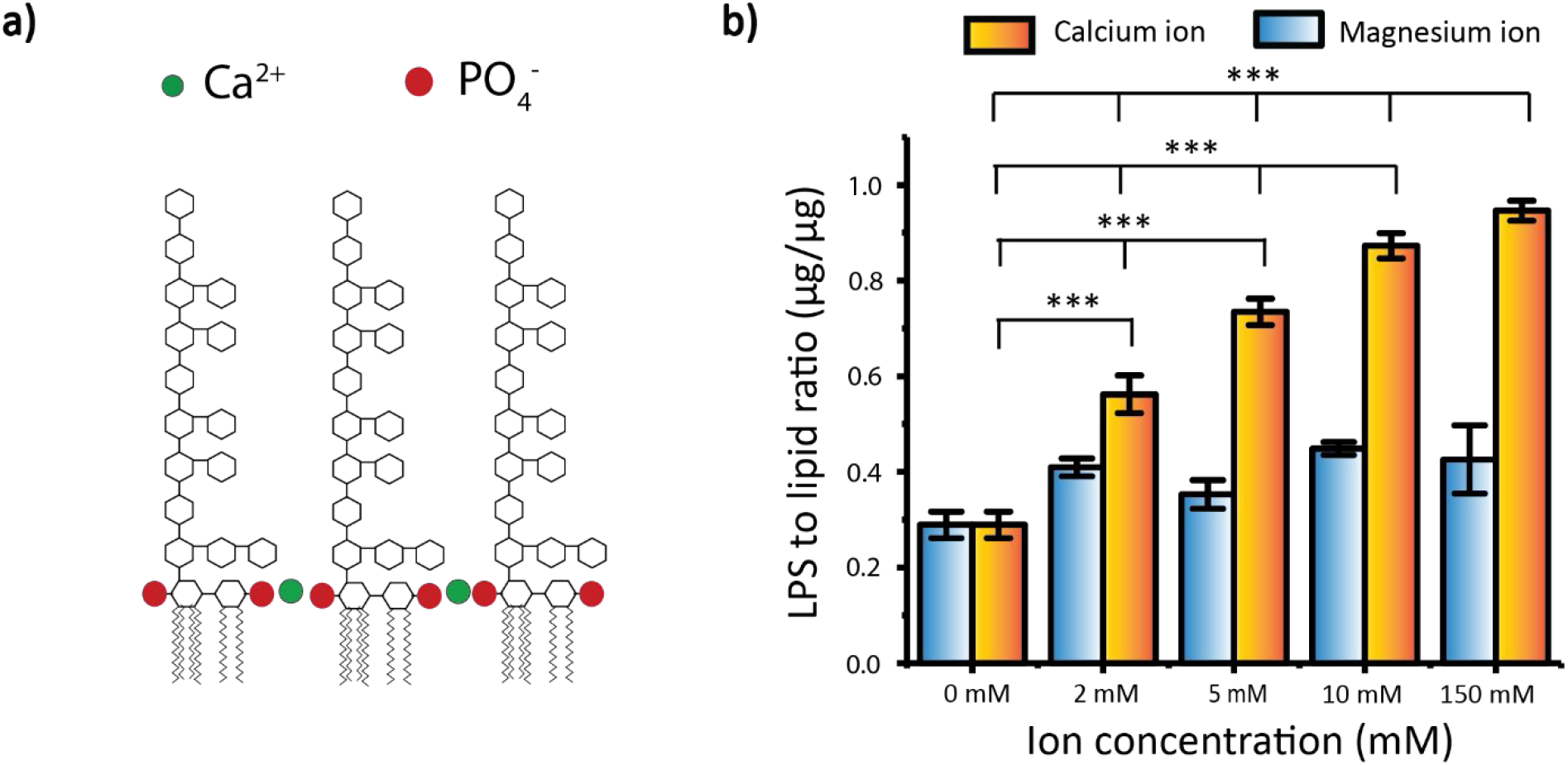
Effect of LPS adsorption by lipid droplets in presence of bivalent ions in the buffer, a) The mechanism proposed by the literature.^29^ A Ca^2+^ ion stays between LPS molecules, stabilizes negatively charged LPS, and promotes self-insertion of LPS. b) Effect of different Mg^2+^ and Ca^2+^ concentrations on LPS adsorption. For analysis, the data was collected from 3 technical replicates for each run. ***p value<0.001, **p value <0.01.

## 3. Discussion

The nature of LPS uptake by lipoprotein plays an essential role in endotoxin removal. In this study, lipid droplets containing phospholipids, triglycerides, and cholesterols but lacking apoproteins were used to mimic the lipoproteins. Fluorescence microscopy results suggested that the amphiphilic LPS stayed majorly on the surface of lipid droplets (**Figure 1**). Comparing the size of lipid droplets using DLS, an increase in the droplet size was observed after LPS uptake (**Table S1**). The DLS data corroborates that LPS is adsorbed on the surface of lipid droplets. The literature suggests that LPS could spontaneously insert into lipid membrane by burying the hydrophobic tail of LPS in the hydrophobic core of lipid bilayer.^35^ We hypothesized that LPS also partitions into the phospholipid monolayer on the surface of the lipid droplets by inserting the hydrophobic tails into the triglyceride cores. Therefore, the effect of LPS adsorption could be influenced by droplet curvature, the surface characteristics of lipid droplets, and the local ionic environment.

According to the literature, the insertion of amphiphilic molecules into cell membranes is curvature sensitive.^18^ Liposomes with high curvatures could introduce defects in phospholipid bilayers, offering the recruitment sites for amphiphilic molecules. Thus, we examined the influence of droplet size on LPS uptake. However, in our experimental condition for lipid droplets (diameter between 70 nm and 500 nm) size does not affect LPS adsorption significantly (**Figure 2a**). Our result is consistent with the other study that evaluated the insertion of fluorescein labeled DHPE into 50–700 nm diameter liposomes.^19^ The impact of liposome size on DHPE-fluorescein insertion was not significant when the liposome size is above 100 nm. For lipoproteins, the curvature dependence of interfacial tension becomes significant when the size of droplets is around 5 nm in radius, if the cross-area of lipid molecules is 1.4 nm^2^.^22^ Because the area per LPS molecule is 1.09 nm^2^,^23^ we expect the curvature may play an important role only in lipid droplets smaller than 10 nm.

The surface molecules of lipid droplets also influence LPS adsorption, including cholesterol and phospholipids. The prior study has shown that cholesterol increases the rigidity of the lipid bilayers and restricts bilayer deformations.^36^ The increase of cholesterol may decelerate the insertion of LPS into lipid droplet surfaces. We observed the increase in cholesterol contents decreased LPS adsorption, probably due to the restriction of LPS insertion into lipid droplet surfaces. The charged phospholipids also affect LPS adsorption. Due to the negatively charged nature of LPS, the increases of the positively charged lipids, such as EPC and DOTAP, enhance LPS adsorption.

It is known that LBP can dissemble LPS aggregates and deliver the LPS monomer to lipoproteins for LPS clearance.^16, 37^ It has been hypothesized that, as a part of the immune response, in sepsis, the human body promotes the production of LBP to aid the removal of LPS by lipoprotein, which reduces the amount of LPS in circulation.^38^ The LBP concentration in the blood for healthy individuals is around 4.1 ± 1.65 µg/ml for healthy individuals and ∼ 31.2 µg/ml (interquartile range, 22.5-47.7 µg /mL) in septic patients.^28^ We also observe LBP promoted LPS adsorption by lipid droplets. The LPS adsorption increase by ∼40% when LBP concentrations increase from 10 µg/ml to 50 µg/ml. This implies LBP could facilitate LPS uptake by lipoprotein. However, LBP can also carry LPS to lymphocyte membranes and trigger TLR4 signaling.^39^ The literature has suggested that LPS neutralization probably depends on the relative amounts of LBP, lipoproteins, signaling proteins, and target cells.^16^

Cations may also affect the electrostatic interactions of LPS on lipid droplets. We observed the increase of Ca^2+^ significantly promote LPS uptake (**Figure 3**) near the physiological concentration. The literature has proposed that Ca^2+^ ions can bridge LPS molecules, stabilize negatively charged LPS, and promote self-insertion of LPS into lipid layers.^29^ However, the LPS uptake rate was not associated with Mg^2+^ at the same concentrations. A similar phenomenon has been observed in other biomolecular interactions involving carboxylate and phosphate groups.^40-42^ The literature suggests that the stability of hydration shells around cations and the polarization effect are the potential factors influencing binding affinities of cations.^42-44^ In contrast to Mg^2+^, Ca^2+^ prefers to form direct contacts with anions, such as carboxylate/phosphate.^43^ This may explain our observations that Ca^2+^ can enhance LPS uptake by directly binding to the negatively charged LPS and bridging them together.

The findings from this study can potentially impact the design of treatment strategies for sepsis. It is known that septic shock patients typically suffer from hypocalcemia (lower level of calcium) and lower levels of lipoproteins. Therefore, the supplementation of either calcium or synthetic lipoprotein has been considered for potential treatments for septic shock patients. However, the preclinical and clinical results are inconclusive.

In the case of calcium supplementation, because the severity and survival rates of septic shock patients were correlated with low Ca^2+^ ion in the blood,^30-32^ calcium replenishment has been recommended to avoid potentially fatal consequences, such as laryngospasm, tetany, seizures, and heart irregularities.^45^ However, no benefit of calcium supplements was found in both preclinical and clinical trials.^46, 47^

In the case of lipoprotein supplementation, phospholipid emulsions have been administrated intravenously to assist in LPS removal. Preclinical studies with injection of endotoxin from *E. coli* followed by the administration of phospholipid emulsions in healthy horses and pigs has shown positive results, as the phospholipid emulsion treatment improves survival in sepsis.^9, 11^ A double-blind, placebo-controlled clinical trial also showed a positive result with emulsion treatment. In this clinical study^8^, before the injection of endotoxin, healthy human volunteers received infusion of emulsion or placebo over a 6-hour period, followed by intravenous administration of endotoxin after two hours. The volunteers who received emulsion had better clinical outcomes (clinical score, temperature, pulse rate) and lower immune response (neutrophil count, tumor necrosis factor–α, and interleukin-6). In another large-scale clinical trial, phospholipid emulsion (850 mg/kg or 1350 mg/kg) and placebo were administered to patients that had confirmed (or were suspected of) gram-negative sepsis. However, there was no reduction in the 28-day mortality or reduction of new organ failure in patients.^7^

The discrepancy between preclinical and clinical findings may be explained by differences in Ca^2+^ concentration. In the small-scale clinical study, the emulsion was given to healthy individuals prior to the injection of LPS.^8^ Although this study did not report the participants’ Ca^2+^ concentrations, their Ca^2+^ concentrations are likely in the normal range as the study used healthy individuals. Therefore, the supplementation of the emulsion probably assisted in the LPS removal. For the large-scale clinical trial^7^, the emulsion was given to the septic patients who typically suffered from hypocalcemia^31^; thus, the low Ca^2+^ concentration in blood may have reduced the efficiency of LPS removal by emulsions. Thus, a combination of calcium and lipid emulsion supplements has promise as a better therapeutic approach. Further clinical studies are needed to test the efficacy of this therapeutic approach against sepsis.

## 4. Conclusion

In summary, we demonstrate that LPS uptake by lipid droplets is controlled by physicochemical properties and environmental factors, such as surface charge, surrounding ions, cholesterol levels, and LBP concentrations. Most interestingly, the LPS uptake rate is significantly influenced by Ca^2+^ ions near physiological concentrations. Because septic shock patients typically suffer from hypocalcemia (lower level of calcium) and lower levels of lipoproteins, the supplementation of Ca^2+^ ions along with synthetic lipoproteins may be a potential therapeutic approach for gram-negative sepsis.

## 5. Methods

### 5.1 Materials

POPC (1-palmitoyl-2-oleoyl-sn-glycero-3-phosphocholine), DOTAP (1,2-Dioleoyl-3-trimethylammonium propane), EPC (1,2-dioleoyl-sn-glycero-3-ethylphosphocholine), POPA (1-palmitoyl-2-oleoyl-sn-glycero-3-phosphate (sodium salt)), POPS (1-palmitoyl-2-oleoyl-sn-glycero-3-phospho-L-serine (sodium salt)) and biotinyl PE (1,2-dioleoyl-sn-glycero-3-phosphoethanolamine-N-(biotinyl) (sodium salt)) were sourced from Avanti polar lipids. Cholesterol (3β-Hydroxy-5-cholestene, 5-Cholesten-3β-ol), glyceryl trioctanoate, LPS-FITC (E. coli serotype O55:B5, CDC 1644-70 strain conjugated to fluorescein isothiocyanate), streptavidin, blocker casein, phosphate-buffered saline (PBS), magnesium chloride, calcium chloride, and HEPES (4-(2-hydroxyethyl)-1-piperazineethanesulfonic acid) were purchased from Millipore Sigma. Texas Red-DHPE (1,2-Dihexadecanoyl-sn-Glycero-3-Phosphoethanolamine, Triethylammonium Salt), Dynabeads Streptavidin T1 were purchased from Fisher Scientific.

### 5.2 Lipid droplet preparation

Lipid droplets were synthesized using the ultrasonic emulsification technique.^48^ Phospholipids, Texas Red-DHPE, cholesterol, and biotinylated lipid, dissolved in chloroform were mixed at the desired composition in a 25 ml glass reactor. Chloroform was then evaporated using a rotary evaporator at 40°C. The lipid thin film was hydrated using 1x PBS solution. After hydration, glyceryl trioctanoate (oil phase) was mixed with the aqueous solution to reach a concentration of 2.5% vol./vol. Subsequently, the sample was sonicated for 15 minutes using a tip sonicator (Qsonica, Q125). During sonication, the samples were kept in an ice bath to prevent overheating. After the tip sonication, the size and zeta potential of the prepared lipid droplets were characterized and stored for further use. The final total lipid concentration was 5 mg/ml.

### 5.3 Size and Zeta potential measurement

The size and zeta potential of lipid droplets were characterized using dynamic light scattering (DLS, Malvern Zetasizer Nano ZS90, Malvern Instruments). A serial dilution of 5 mg/ml lipid droplet solution was prepared to minimize multi-particle scattering. The size measurement remained constant irrespective of the dilution factor. Zeta potential was measured at room temperature using disposable folded capillary cells. A minimum of three measurements per sample were taken.

### 5.4 LPS adsorption measured by fluorescent spectroscopy

All the experiments were carried out at a physiological pH of 7.4. For this purpose, 1xPBS buffer was used except for the study of bivalent ions where 20 mM HEPES buffer was used. In all the experiments, the total lipid amount was kept constant. The stock solution of lipid droplets (5 mg/ml total lipid) was diluted 100-fold with the desired buffers. Then, 100 µl of diluted lipid droplets, 24 µl streptavidin coated Dynabeads, and the desired volume of buffer (1x PBS) with or without LBP were mixed. In the study of bivalent ions, all the dilutions and pH were maintained using 20 mM HEPES buffer. CaCl_2_ or MgCl_2_ salt solution in 20 mM HEPES buffer (final pH 7.4) was added to get the final concentration from 0 to 150 mM. The mixture was incubated at 37 °C for 2 hours. Streptavidin on Dynabeads would capture lipid droplets decorated with biotinylated lipids. The lipid droplet solutions were then incubated with 100 µl of 1 mg/ml of the LPS labeled with the fluorophores (FITC-LPS) for 2 hours at 37 °C. Dynabeads were then separated from the solutions using a magnetic rack. The supernatants of lipid droplet-LPS mixtures were collected, and the fluorescent signal was measured using the FLUOstar Omega Microplate Reader (**Figure 4**).

**Figure 4.**
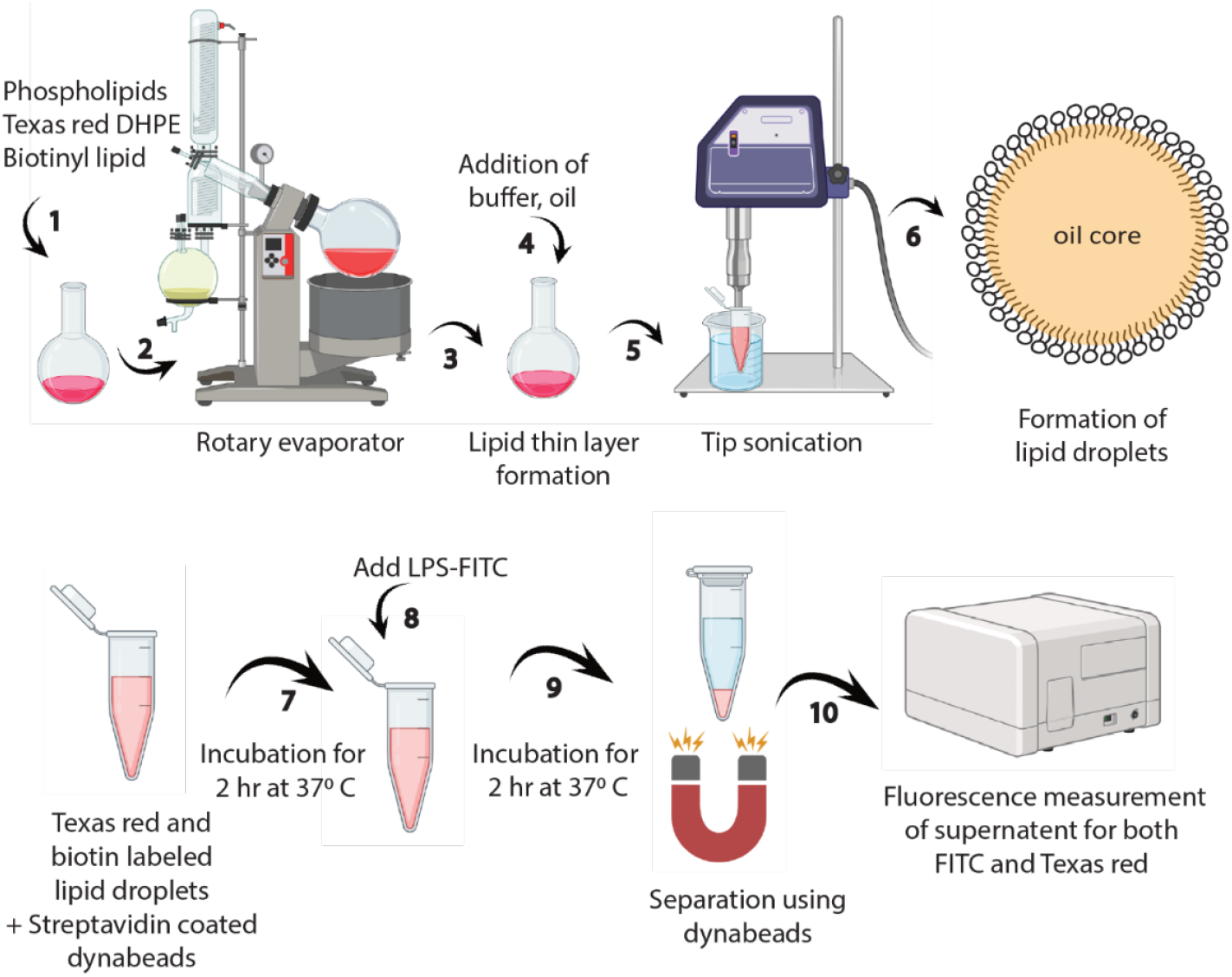
Schematic for lipid droplet preparation and studying their interactions with lipopolysaccharides.

The LPS uptake by lipid droplets was quantified by the fluorescence signals. The calibration samples containing the known concentrations of FITC-LPS, and Texas Red labeled droplets were prepared along with the uptake experiments. The amounts of LPS uptakes by the lipid droplets that were captured by Dynabeads were calculated by comparing them with the calibration samples. Because the total number of surface lipids would influence the capacity of LPS adsorption, the LPS uptake rates were reported as the mass ratios of adsorbed LPS to lipid molecules.

### 5.5 Imaging LPS adsorption on giant lipid droplets using epifluorescence microscopy

POPC, Texas Red-DHPE, and biotinylated lipid were mixed at the desired composition in a glass reactor, and chloroform was evaporated using a rotary evaporator. The lipid thin film was then hydrated using double deionized (DDI) water. The lipid solution was mixed with glyceryl trioctanoate in a 5:1 volume ratio. The mixture was vortexed for 3 minutes and diluted 10-fold with DDI water. Giant lipid droplets were then separated from smaller lipid droplets using a centrifuge at 1000 RCF for 5 minutes. A 96-well flat bottom plate coated with streptavidin was prepared for capturing lipid droplets. The well plate with a glass bottom was etched with 1N sulfuric acid, rinsed with DDI water, and then incubated with the streptavidin solution. The well plate was then blocked with 0.1% casein solution for 1 hour. Giant lipid droplets were adsorbed on glass surfaces via streptavidin-biotin bonding. The unbound droplets were washed with the desired buffer. FITC-LPS was added to the well plate and incubated for 2 hours. The excess FITC-LPS was removed by washing the wells. The adsorption of LPS on giant lipid droplets was observed using Zeiss Axiovert 200M fluorescence microscope with a 20x objective. Giant lipid droplets labeled with Texas Red-DHPE were observed by the Texas Red channel (absorption wavelength 585 nm, emission wavelength 620 nm) and LPS adsorptions were observed by the FITC channel (absorption wavelength 485 nm, emission wavelength 520 nm). The images were processed using ImageJ software.

## AUTHOR INFORMATION

### Author Contributions

The manuscript was written through contributions of all authors.

## ACKNOWLEDGMENT

The authors gratefully thank Professor Jeetain Mittal at Texas A&M University for insightful discussion. The authors acknowledge the support from National Institutes of Health (R03AI139650).

Parts of Figure 4 were created with BioRender (Arnob, A. (2024) BioRender.com/y86q069).

## ABBREVIATIONS

DHPE: 1,2-Dihexadecanoyl-*sn*-Glycero-3-Phosphoethanolamine, Triethylammonium Salt
HEPES: 4-(2-hydroxyethyl)-1-piperazineethanesulfonic acid
LPS: lipopolysaccharide
LBP: lipopolysaccharide binding protein
FITC: Fluorescein isothiocyanate
FITC-LPS: Fluorescein isothiocyanate (FITC) labeled lipopolysaccharide (LPS)
VLDL: very low-density lipoproteins
LDL: low-density lipoproteins
IDL: Intermediate-density lipoproteins
HDL: high-density lipoproteins
POPC: 1-palmitoyl-2-oleoyl-sn-glycero-3-phosphocholine
DOTAP: 1,2-Dioleoyl-3-trimethylammonium propane
EPC: 1,2-dioleoyl-sn-glycero-3-ethylphosphocholine
PA: Phosphatidic Acid
PS: Phosphatidylserine

## Notes

### Competing Interest Statement

The authors have declared no competing interest.

