## Supplementary material for "Factors Promoting Lipopolysaccharide Uptake by Synthetic Lipid Droplets"

**Table S1: DLS size and zeta potential measurement of lipid droplets in 1x PBS buffer**

| Lipid droplet compositions | Before LPS adsorption | | After LPS adsorption | |
|---|---|---|---|---|
| | size | zeta potential | size | zeta potential |
| 99.25 % POPC | 211.6 ± 0.9 | -7.5 ± 0.5 | 223.3 ± 2.8 | -4.5 ± 0.1 |
| 10 % DOTAP + 89.25 % POPC | 202.3 ± 0.5 | 9.3 ± 0.3 | 213.6 ± 0.1 | 1.4 ± 0.1 |
| 10 % EPC + 89.25 % POPC | 196.8 ± .03 | 6.7 ± 0.2 | 210.3 ± 0.2 | 0.0 ± 0.3 |
| 20 % POPS + 79.25 % POPC | 199.9 ± 0.2 | -17.8 ± 0.5 | 211.5 ± 0.2 | -11.1 ± 0.3 |
| 5 % POPA + 94.25 % POPC | 202.6 ± 0.3 | -6.7 ± 0.9 | 213.5 ± 0.4 | -5.0 ± 0.4 |

**Table S2: Molar composition of lipid droplets for characterization and LPS adsorption study.**

| **Study parameter** | | **POPC (mol %)** | **POPA (mol%)** | **POPS (mol%)** | **DOTAP (mol%)** | **EPC (mol%)** | **Cholesterol (mol%)** | **Texas Red (mol%)** | **Biotin (mol%)** | **Triglycerides (µl)** |
|---|---|---|---|---|---|---|---|---|---|---|
| Size effect | 70.3 ± 0.9 nm | 99.25 | 0 | 0 | 0 | 0 | 0 | 0.25 | 0.5 | 2 |
| | 219.2 ± 0.3 nm | 99.25 | 0 | 0 | 0 | 0 | 0 | 0.25 | 0.5 | 20 |
| | 484.7 ± 0.5 nm | 99.25 | 0 | 0 | 0 | 0 | 0 | 0.25 | 0.5 | 40 |
| Cholesterol effect | 0 mol% | 99.25 | 0 | 0 | 0 | 0 | 0 | 0.25 | 0.5 | 20 |
| | 10 mol% | 89.25 | 0 | 0 | 0 | 0 | 10 | 0.25 | 0.5 | 20 |
| | 20 mol% | 79.25 | 0 | 0 | 0 | 0 | 20 | 0.25 | 0.5 | 20 |
| | 30 mol% | 69.25 | 0 | 0 | 0 | 0 | 30 | 0.25 | 0.5 | 20 |
| Charge effect | Positive charge | 89.25 | 0 | 0 | 0 | 10 | 0 | 0.25 | 0.5 | 20 |
| | | 89.25 | 0 | 0 | 10 | 0 | 0 | 0.25 | 0.5 | 20 |
| | Negative charge | 79.25 | 0 | 20 | 0 | 0 | 0 | 0.25 | 0.5 | 20 |
| | | 94.25 | 5 | 0 | 0 | 0 | 0 | 0.25 | 0.5 | 20 |

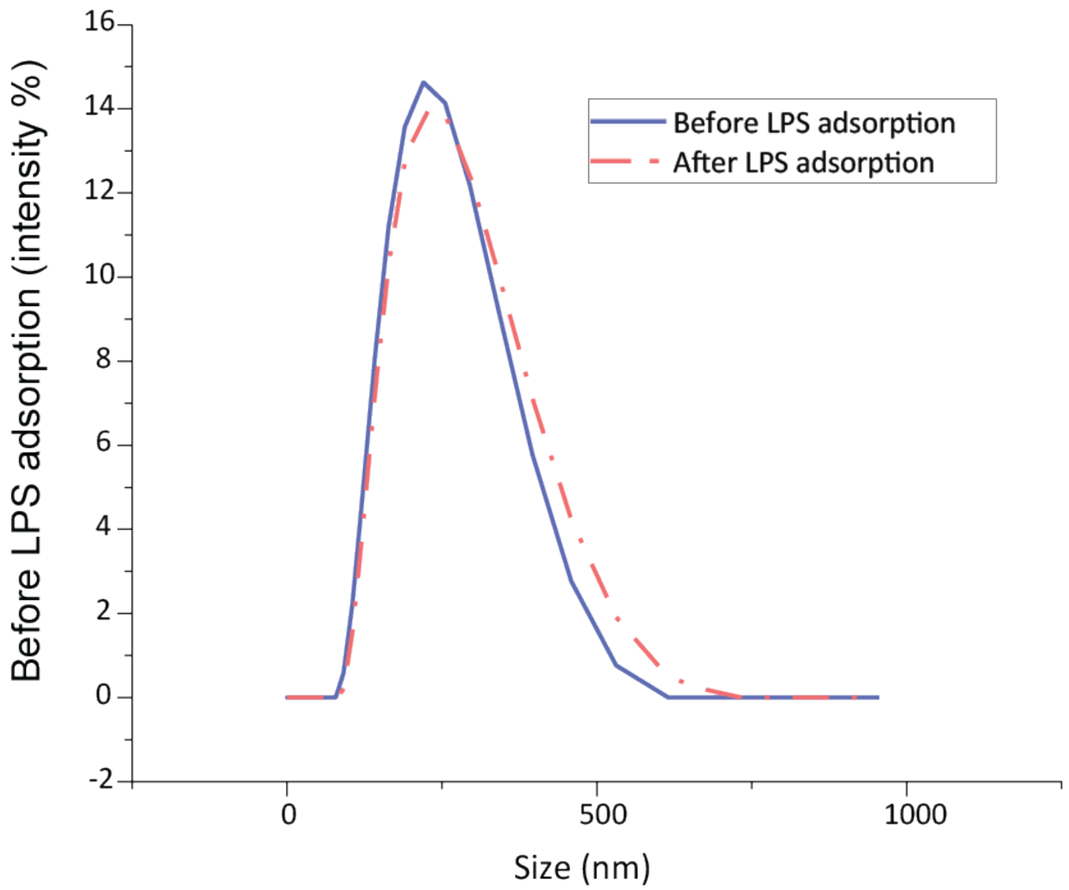


**Figure S1. DLS size measurement (average, n=3) of lipid droplet (99.25% POPC) before and after LPS adsorption at $25^0$ C for 2hr.**
